# Linking animal migration and ecosystem processes: data-driven simulation of propagule dispersal by migratory herbivores

**DOI:** 10.1101/2021.05.21.445111

**Authors:** Marius Somveille, Diego Ellis-Soto

## Abstract

1. Animal migration is a key process underlying active subsidies and species dispersal over long distances, which affects the connectivity and functioning of ecosystems. Despite much research describing patterns of where animals migrate, we still lack a framework for quantifying and predicting how animal migration affects ecosystem processes.
2. In this study, we aim to integrate animal movement behavior and ecosystem functioning by developing a predictive modeling framework that can inform ecosystem management and conservation. Our framework models individual-level migration trajectories between populations’ seasonal ranges as well as the resulting dispersal and fate of propagules carried by the migratory animals, and it can be calibrated using empirical data at every step of the modeling process.
3. As a case study, we applied our framework to model the spread of guava seeds, *Psidium guajava*, by a population of migratory Galapagos tortoises, *Chelonoidis porteri*, across Santa Cruz Island. Galapagos tortoises are large herbivores that transport seeds and nutrients across the island, while Guava is one of the most problematic invasive species in the Galapagos archipelago.
4. Our model is able to predict the pattern of spread of guava seeds alongside tortoises’ downslope migration range, and it identified areas most likely to see germination success and establishment. Our results show that Galapagos tortoises’ seed dispersal may particularly contribute to guava range expansion on Santa Cruz Island, due to both long gut retention time and tortoise’s long-distance migration across vegetation zones. In particular, we predict that tortoises are dispersing a significant amount of guava seeds into the Galapagos National Park, which has important consequences for the native flora.
5. The flexibility and modularity of our framework allows for the integration of multiple data sources. It also allows for a wide range of applications to investigate how migratory animals affect ecosystem processes, including propagule dispersal but also other processes such as nutrient transport across ecosystems. Our framework is also a valuable tool for predicting how animal-mediated propagule dispersal can be affected by environmental change. These different applications can have important conservation implications for the management of ecosystems that include migratory animals.

## 1.0 Introduction

Ecosystems are connected through flows of energy and materials transported passively by the abiotic environment or actively through the transport of individuals, gametes or spores (Loreau *et al*. 2003). The movement of animals is a key process shaping active subsidies and dispersal of plants, which affects the functioning and connectivity of ecosystems (Côrtes & Uriarte 2013; Earl & Zollner 2017; Schmitz *et al*. 2018; Subalusky & Post 2018; Ellis-Soto *et al*. 2020). In particular, animal migration – the regular, directional movement of animals between specific destinations – involves billions of animals across the planet (Hu *et al*. 2016; Dokter *et al*. 2018) and provides important ecosystem services (Bauer & Hoye 2014). However, despite the ecological importance of the migration phenomenon and the research efforts to describe patterns of where animals migrate, we still lack a predictive framework that we can use to quantify how migratory animals impact ecosystem functioning.

One of the main ecosystem services provided by migratory animals is propagule dispersal, and particularly seeds (Bauer & Hoye 2014). More than half of all plant species are dispersed by animals (Aslan *et al*. 2013), and such animal-mediated seed dispersal influences plant species survival and range expansion into new environments (Nathan & Muller-Landau 2000; Kendrick *et al*. 2012; Travis *et al*. 2013). The movement ecology of seeds has thus been highlighted as a key knowledge gap to improve our understanding of plant distribution (Beckman *et al*. 2019). Quantifying and predicting the role of animals in seed dispersal is also particularly relevant given that the scale of animal movements is declining globally in response to human activities (Tucker *et al*. 2021), which has important implications for plant-animal interactions (Neuschulz *et al*. 2016). Animal migrations, in particular, are disappearing at alarming rates (Wilcove & Wikelski 2008) with unknown, but likely significant, consequences on ecosystem functioning. For instance, the loss of Pleistocene megafauna and their long ranging movements is thought to have significantly reduced seed dispersal across large spatial scales (Malhi *et al*. 2016; Pires *et al*. 2018).

Dispersal kernels, which allow modeling how far seeds can be dispersed, have been usually employed to understand dispersal of plant species (Nathan 2006; Nathan *et al*. 2012; Bullock *et al*. 2017; Pires *et al*. 2018). For animal-mediated seed dispersal, however, it is important to take into account intra-specific variation in the seed dispersal ability of animals (Zwolak 2018). In addition, modeling seed dispersal must ideally be spatially-explicit in order to account for the context dependency of where seeds are deposited and the probability of successful germination and establishment (Nathan & Muller-Landau 2000). Developments in tracking technologies allow quantification of the movement of animals at fine spatio-temporal scales for long periods of time (Kays *et al*. 2015). This opens up opportunities to study animal movement and understand its underlying environmental and internal drivers (Nathan *et al*.2008; Hawkes *et al*. 2011; Jesmer *et al*. 2018). Recent years have seen an increase in studies coupling GPS tracking of animals with ecosystem processes, especially seed dispersal (Kleyheeg *et al*. 2017, 2019; Oleksy *et al*. 2017; van Toor *et al*. 2019). This approach provides a spatially explicit, individual-based understanding of seed dispersal by migratory animals that integrates animal movement, seeds dispersed and gut retention times. However, previous studies have only modeled dispersal from a single location, which limits the potential applications of such approach. In addition, these studies do not use empirical data explicitly to model the fate of the dispersed seeds once released in the environment.

Here we propose a novel modeling framework that combines simulation and empirical data from a variety of sources to model the spread of seeds by migratory herbivores. Our framework allows to model individual-level migration trajectories between the seasonal distributions of populations together with the resulting dispersal and fate of seeds carried by the migratory animals, and it allows using empirical data to calibrate every steps of the modeling process. This approach aims to quantitatively harmonize behavioral ecology and ecosystem process and make quantitative predictions that could inform ecosystem management and conservation. As a case study, we applied our framework to model the spread of guava seeds, *Psidium guajava* (Linnaeus), by a population of migratory Galapagos tortoises, *Chelonoidis porteri* (Rothschild) (IUCN critically endangered), across Santa Cruz Island. Galapagos tortoises are the largest terrestrial ectotherms worldwide and are considered ecosystem engineers through the transport of seeds and nutrients, as well as herbivory and trampling (Gibbs *et al*. 2010; Blake *et al*. 2012; Ellis-Soto 2020), while Guava is one of the most problematic invasive species in the Galapagos archipelago, having been introduced in the late 19^th^ century (Walsh *et al*. 2008).

Santa Cruz Island contains the highest human population on the Galapagos. The island harbors more introduced plant species that native species (Guézou *et al*. 2010), which threatens endemic biodiversity. On Santa Cruz Island, presence of invasive guava has modified native plant community and ecosystems, especially so in the humid highlands (Guézou *et al*. 2010), and several ongoing eradication initiatives have so far proven unsuccessful (Gardener *et al*.2010). Guava is heavily consumed by Galapagos tortoises (Blake *et al*. 2015), which perform seasonal migrations across the elevational gradient of the island, and disperse guava seeds throughout their range (Blake *et al*. 2012, 2013; Ellis-Soto *et al*. 2017). Most of the island’s surface is located within the Galapagos National Park (GNP) and surrounds heavily degraded agricultural land where invasive species such as guava are widespread (Trueman *et al*. 2014; Benitez-Capistros *et al*. 2019). The permeability of national park boundary to agricultural lands means that animals like tortoises can regularly cross from the park into farm land, where they consume introduced plants including fruits, before returning to the park (Benitez-Capistros *et al*. 2018). Taking advantage of a long-term study on Galapagos tortoises (Blake *et al*. 2012, 2013, 2015; Yackulic *et al*. 2016; Benitez-Capistros *et al*. 2019, Sadeghayobi *et al*. 2011), we use data on tortoise ecology, migratory behavior, seed dispersal ability, diet preference and gut retention times in order to calibrate our modeling framework and simulate how tortoises spread guava across Santa Cruz island. Specifically, we aim to (i) simulate realistic migration trajectories for tortoises across the population’s geographical range, (ii) model how guava seeds are spread by migrating tortoises, and (iii) use empirical data to calibrate the model and validate the seed dispersal predictions.

## 2.0 Material and Methods

### 2.1 Empirical data

#### 2.1.1 Study site and habitat

We obtained a landcover map for Santa Cruz Island, Galapagos, from (Rivas-Torres *et al*. 2018a) and shapefiles of the Galapagos National Park and agricultural land from the 2014 census conducted by the Ecuadorian Ministry of Agriculture (CGREG 2015). This allowed us to estimate the proportion of different habitats where invasive guava is deposited through seed dispersal by *C. porteri* as well as the ratio of seed deposition in national park and agricultural land to better understand the context dependency of these events. To estimate available resources for tortoises, we used the Normalized Difference Vegetation Index (NDVI), a remote-sensing measure of greenness that correlates well with primary productivity, and which has been shown to be an important drivers of tortoises annual migration (Blake *et al*. 2013; Yackulic *et al*. 2016; Bastille-Rousseau *et al*. 2019). To estimate the spatial distribution of guava, we made use of a recently published land cover classification of the Galapagos which provides the distribution of guava patches at very high resolution in Santa Cruz Island (Laso *et al*. 2019).

To estimate whether guava seeds can germinate and become adult reproductive trees, we made use of a species distribution model (SDM) for guava previously published (Ellis-Soto *et al*. 2017). This model relates environmental predictors (i.e., temperature and precipitation long term averages and seasonality commonly referred to as bioclimatic variables; (Hijmans *et al*. 2005)) to georeferenced guava locations and indicates how suitable a given location in Santa Cruz is for this plant species. This model was then overlaid on adult guava trees on Santa Cruz to understand at which suitability score guava plants can germinate, survive and establish under current and future climatic conditions.

#### 2.1.2 Tortoise migratory movements

We used tracking data of 9 tagged adult male Galapagos giant tortoises for which hourly locations were recorded between 2013 and 2018 (attachment procedures and GPS sampling regimes are described in Blake *et al*. 2013). To identify the starting and end locations of individual downslope migrations between 2013 and 2018, we made use of the locator() function from the graphics package in R on plots representing individual tortoise net square displacement (Singh *et al*. 2016), which provided us with 16,464 GPS tortoise locations. We removed empirical migratory tracks that were too sinuous as their characteristics generate convergence issues for the model described below. We retained 19 downslope migration trajectories from 5 individuals (i.e. individuals were tracked for several years; Fig. S1), sampling each track to one point per day (i.e., taking the first GPS point when multiple points where present for a single day).

#### 2.1.3 Tortoise diet and gut retention

Sampling of 222 tortoise dung piles revealed that guava is the most common dispersed plant species by *C. porteri* with an average of 624 seeds per dung pile (Ellis-Soto *et al*. 2017). Feeding trial experiments with pseudo seeds identified the gut retention time of *C. porteri* during the period of downslope migration (*μ* = 7.5, σ = 2.16 in days, respectively) (Sadeghayobi *et al*. 2011), which applies to guava seeds. In addition, *ex situ* germination trials suggests that tortoise ingestion and dung does not influence germination success of guava seeds neither positively nor negatively (Blake *et al*. 2012).

### 2.2 Modeling framework

#### 2.2.1 Model overview

We simulated the migratory movement of tortoises from the highlands to the lowlands of Santa Cruz Island using a two-step modeling process: first, we simulated the migratory connectivity of the population (i.e. links between seasonally-occupied sites based on individual tortoises migratory movements), and second, we simulated the trajectory of migrating individuals (Fig. 1).

**Figure 1.**
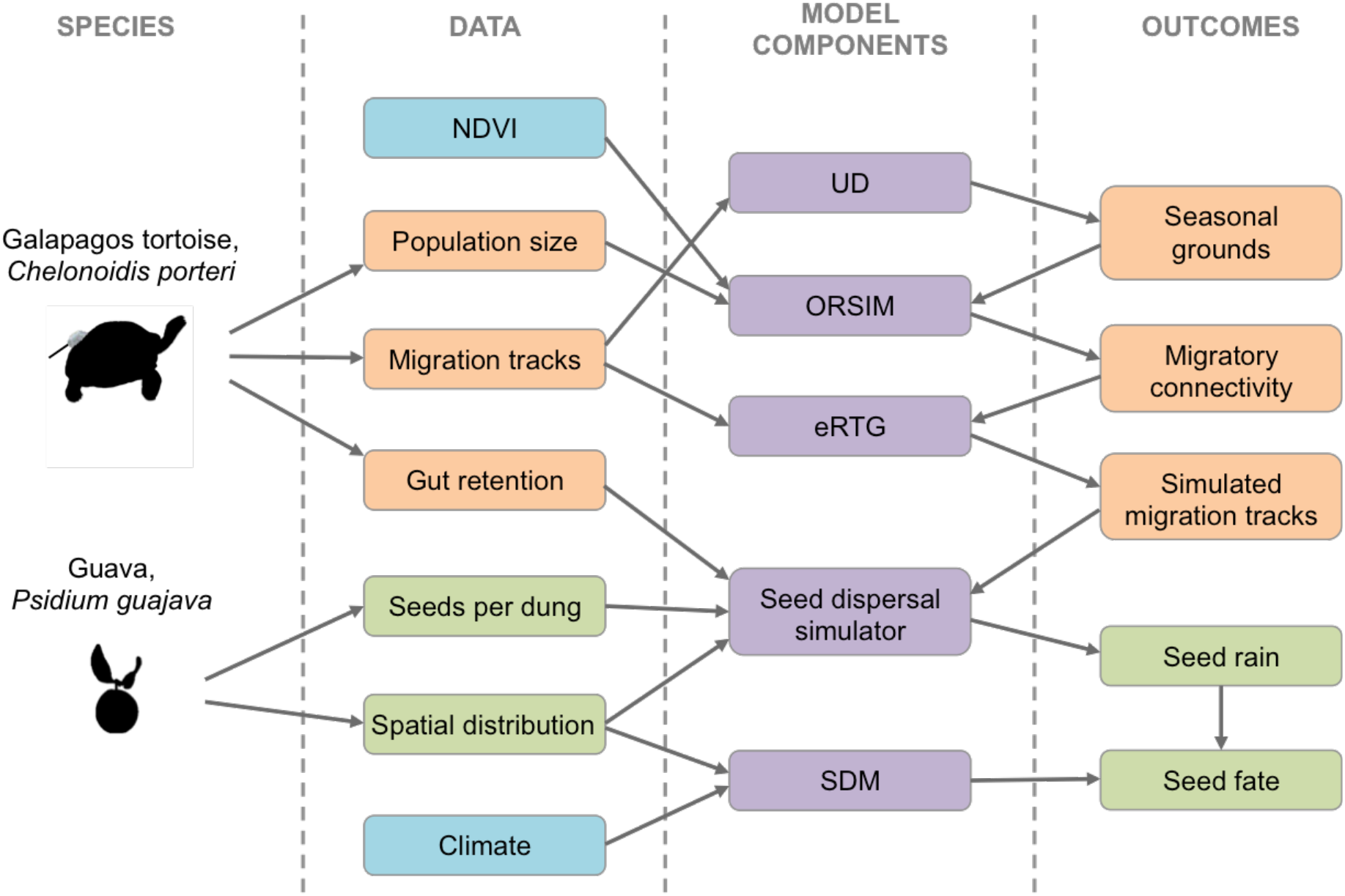
Workflow for data-driven simulation of seed dispersal by a migrating species. Orange boxes indicate data and simulation outcomes for migratory Galapagos tortoises; green boxes indicate data and simulation outcomes for the invasive plant species guava; blue boxes indicate environmental data; and purple boxes indicate model components (see Material and Methods for details). NDVI: normalized difference vegetation index, a remote sensing measure capturing primary productivity; UD: utilization distributions, computed from empirical movement tracks; ORSIM: optimal redistribution simulator, which simulates migratory connectivity; eRTG: empirical random trajectory generator, which generates a movement between two end points that is calibrated by empirical migration tracks; SDM: species distribution model, indicating guava germination success. Seed rain refers to the spatial distribution of the density of seeds resulting from seed dispersal by migrating tortoises; seed fate refers to the spatial distribution of the density of germinated seeds resulting from seed dispersal by migrating tortoises.

#### 2.2.2 Simulating migratory connectivity

We simulated migratory connectivity between the seasonal distributions of tortoises, i.e., in the highlands and in the lowlands. To map areas seasonally utilized by tortoises, we calculated 95 percentile utilization distribution (UD) (Fieberg & Kochanny 2005) for each of the nine tortoises (see 2.1.2) using the adehabitatHR package (Calenge 2006). We then drew a geometric convex hull around these UD’s to obtain a population-level highland and lowland range. Highland and lowland areas were converted into presences and absences on a grid of hexagons with equal area (~1.18 km^2^) and shape covering Santa Cruz Island.

To simulate migratory connectivity between the seasonal distributions of tortoises, we used a model called the Optimal Redistribution Simulator (ORSIM), which was shown to capture well avian migratory connectivity patterns (Somveille *et al*. in review). This model is based on energy optimization, and captures two processes: minimizing energetic costs associated with relocating between seasonal grounds, and intra-specific competition for access to energy supply. ORSIM uses a solution to the Monge-Kantorovich transportation problem (Hitchcock 1941; Rachev 1984) from linear optimization, which can be formalized linear programming as follows.

Let *H* = {(*h*_1_, *k*_*h*_1__),…,(*h_m_*, *k_h_m__*)} be the distribution of energy available during the season spent in the highland, where *h_i_* is highland site *i* and *k_h_i__* is the weight of this site, which corresponds to the energy available at this site; and *L* = {(*l*_1_, *k*_*l*_1__),…,(*l_n_*, *k_l_n__*)} be the distribution of energy available during the season spent in the lowland, with *n* lowland sites, where *l_j_* is lowland site *j* and *k_l_j__* is the weight of this site, which corresponds to its energy supply. We want to find a total flow *F* = [*f_ij_*], with *f_ij_* the flow of energy between *h_i_* and *l_j_*, that minimizes the overall cost

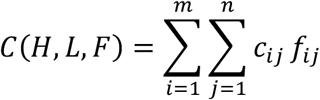

where *C_ij_* is the energetic cost associated with relocating between sites *h_i_* and *l_j_*. This function is subject to the following constraints:

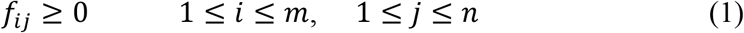

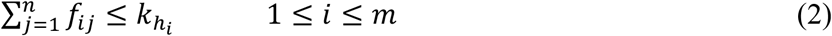

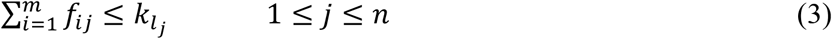

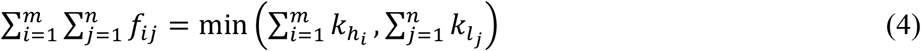

Constraint (1) allows energy to move from *H* to *L* and not vice versa. Constraint (2) limits the amount of energy that can move away from the highland sites in *H* and reflects the energy demand of departing individuals. Constraint (3) limits the lowland sites in *L* to receive no more energy than their energy supply. Finally, constraint (4) specifies that the total amount of energy must be equal to either the total energy demand of highland sites or the total energy supply of lowland sites, whichever one is the smallest, thus forcing to move the maximum amount of energy possible.

This modeling framework assumes that energy is transferred between highland and lowland sites via migrating tortoises, which are energetically equivalent, i.e., they all have the same energetic needs and cost function, and the same competitive ability. Energy availability across the highland and lowland distributions of tortoises was estimated using NDVI values (see 2.1.1). We rescaled these NDVI values so that a total of 1000 migrating individuals were generated by the model. For a complete description of ORSIM including more details about the underlying assumptions see Somveille *et al*. (in revision).

A solution to the transportation problem from linear optimization is implemented in the Earth Mover’s Distance (EMD) algorithm (Rubner *et al*. 2000), which uses the transportation-simplex method (Hillier & Lieberman 1990). To simulate migratory connectivity, we used the FastEMD algorithm (Pele & Werman 2008, 2009), which is implemented in the Python wrapper PyEMD. No distance threshold was used when running FastEMD.

#### 2.2.3 Simulating migration trajectories

We simulated the explicit migration trajectories of migrating tortoises between their starting point in the highland and the respective destinations in the lowland that are generated by the migratory connectivity model (2.2.2). To do so, we used the empirical Random Trajectory Generator (eRTG; Technitis *et al*. 2015) in the R environment. This algorithm generates the movement between two endpoints with a fixed number of steps, which retains the geometric characteristics of real observed trajectories based on tracking data (see detailed description in Kleyheeg et al. 2019 and van Toor et al. 2019). The eRTG is similar to a biased correlated random walk and can be best described as a mean-reverting Ornstein-Uhlenbeck process (Smouse *et al*. 2010). The algorithm uses empirical tracking data as a template, and takes empirical distributions of the following characteristics of animal movement: step length, turning angles, their autocorrelation at a lag of one step, and the covariance of step length and turning angle. We estimated the distributions of these movement characteristics using the empirical tracking data on tortoises’ downslope migratory movement described above.

Using the eRTG, we simulated the movement trajectories of the 1000 migrating tortoises generated by the model of migratory connectivity. For each simulated migrating tortoise, we used the following procedure: we ran eRTG 10 times with the shortest empirical track (i.e., with the least number of movement daily steps); if the model converged, we selected the resulting simulated trajectory, but if the model did not converge at least once, we then ran eRTG 10 times with the second shortest empirical track; and we continued for all 19 empirical tracks until the model converged. All models (i.e., for each simulated individual) ultimately converged.

#### 2.2.4 Simulating seed dispersal

To simulate the spread of guava seeds by migrating tortoises, we combined the simulated migration trajectories with an empirically informed simulator of tortoises’ consumption and excretion of guava seeds. In our model, tortoises are assumed to engage in migratory movement once a day, although we also ran a sensitivity analysis in which tortoises move twice and four times a day. During the rest of the day when they are not migrating, tortoises are assumed to stay put, eat and produce excrements. When not migrating, if a tortoise is located where guava is present, we assumed that it eats guava. In addition, each tortoise has a probability of excreting guava seeds based on gut retention time and the results of previous feeding events, determined as:

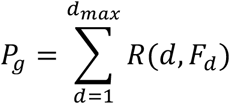

with *P_g_*: probability of excreting guava seeds; *d*: number of days since a feeding event; *F_d_*: results of foraging event at *d* (i.e. whether or not guava was eaten); *R*: gut retention time distribution, which was estimated empirically as a truncated normal distribution with *μ* = 7.5 and σ = 2.16 (Sadeghayobi *et al*. 2011); and *d_max_* indicates a threshold, which was set to 20 days, above which guava seeds were considered to be no longer present in the gut. For each of the 20 days preceding the start of the simulation, a tortoise was set to have eaten guava given probability = 0.21, which corresponds to the proportion of the highland distribution that is covered by the spatial distribution of guava.

To determine whether guava is excreted when a tortoise in not migrating, we sampled a value between 0 (no guava excreted) and 1 (guava excreted) based on *P_g_*. Then, if guava is excreted, the number of guava seeds excreted during the excretion event is determined by randomly sampling an empirically-estimated truncated normal distribution with *μ* = 1443 and σ = 2057 (Ellis-Soto *et al*. 2017). Using this methodology, we modeled where and how many guava seeds are excreted along the migratory trajectory of each simulated migrating tortoises, which we call ‘seed rain’. We then overlaid the seed rain on the projection of the SDM that predict germination success (i.e. binary value of successful versus non-successful germination, based on Ellis-Soto *et al*. 2017; see section 2.1.1) in order to determine whether seeds are likely to germinate or not (i.e. seed fate).

### 2.3 Model validation

To validate the seed dispersal prediction of our model, we used information of collected dung piles containing guava seeds from Ellis-Soto et al. (2017) and added a few others opportunistically collected since them. We obtained a total of 101 dung piles containing guava across an elevation gradient from 28m to 419m. To better understand the spatial context in which tortoise-dispersed guava seeds are deposited across our study area, we associated dung pile occurrences with landcover classes of Santa Cruz Island from (Rivas-Torres *et al*. 2018b). We focused specifically on an area locally known as ‘La Reserva’ to the south and southwest of the island, which is the core distribution area of *C. porteri* (Ellis-Soto et al. 2017). We extracted landcover information at the location of the observed and simulated dung piles containing guava. We inspected whether the frequency density of dung piles containing guava seeds across elevation as well as the distribution of dung piles containing guava seeds across habitats, in particular in agricultural areas versus the Galapagos National Park, were similar for simulated versus observed dung piles.

### 2.4 Seed fate

Besides landcover, we made use of a previously estimated species distribution model (SDM) for guava as a proxy of seed fate for seeds deposited across a gradient of climatic suitability. We used the thresholded guava SDM from Ellis-Soto et al. (2017) to identify suitable habitat for establishment of guava (seed dispersal efficiency). Briefly, the threshold was chosen based on the minimum climatic suitability in which an actual guava plant was observed during a vegetation survey across Santa Cruz Island.

## 3.0 Results

### 3.1 Simulated and observed seed dispersal by migratory tortoises

Our data-driven modeling framework was able to simulate the downslope migration of adult male Galapagos tortoises between highland and lowland seasonal grounds on Santa Cruz Island (Fig. 2a). The vast majority of trajectories and the overall migration pattern of the population simulated by the model appear realistic. The resulting dispersal of guava seeds by migrating tortoises spreads from the highland distribution to the lowland distribution, decreasing in intensity as tortoises arrive closer to the lowlands (Fig. 2b). This pattern matches well empirical data of dung piles containing guava seeds that were not used to parameterize the model (Fig. 3, Fig. S2). The occurrences of observed and simulated dung piles containing guava peak at ca. 180m and 150m in elevation respectively, with a decay towards lower (Galapagos National Park areas) and higher (Agricultural and privately owned areas) elevations (Fig. 3a,c). However, simulated guava seeds appear to be somewhat higher than observed at lower elevations, c.a. 80m, and lower than observed at high elevation, c.a. 400m (Fig. 3a,c, Fig. S2). Both observed and simulated dung piles containing guava seeds are outside the current elevational range of guava in Santa Cruz Island, potentially facilitating the expansion of this invasive species (Fig. 3). The sensitivity analysis of the number of times tortoises engage in migratory movement per day indicates that simulation results match empirical data less well when modeled tortoises were migrating, eating and execrating more often, in particular because it increases unrealistically the long-distance dispersal of guava seeds to the lowland (Fig. S3).

**Figure 2.**
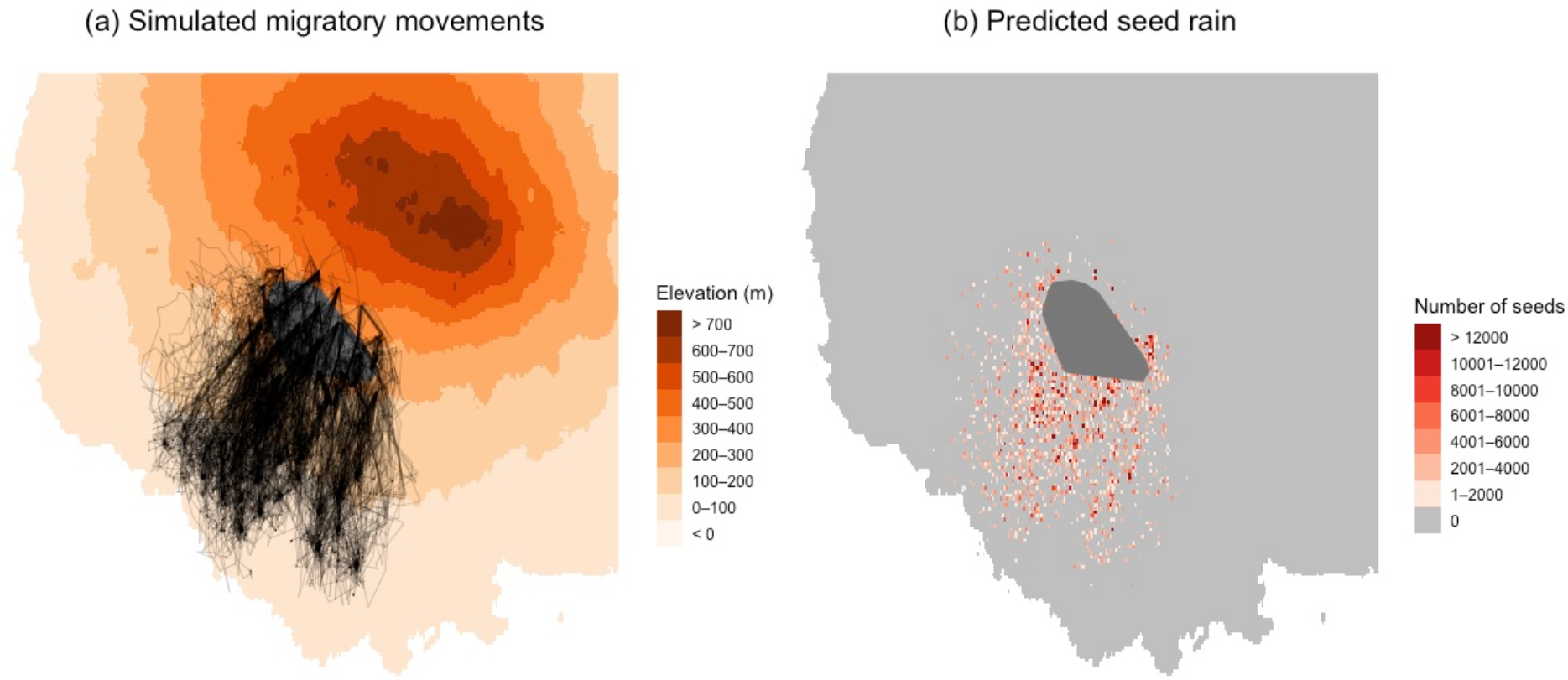
Simulated tortoise migrations and resulting guava seed rain. (a) Simulated migration trajectories of tortoises, from highland to lowland. Each black line indicates the simulated migration of an individual. (b) Density of guava seeds dispersed by simulated migrating tortoises, also called ‘seed rain’. The highland seasonal distribution of the population is represented by the grey polygon.

**Figure 3.**
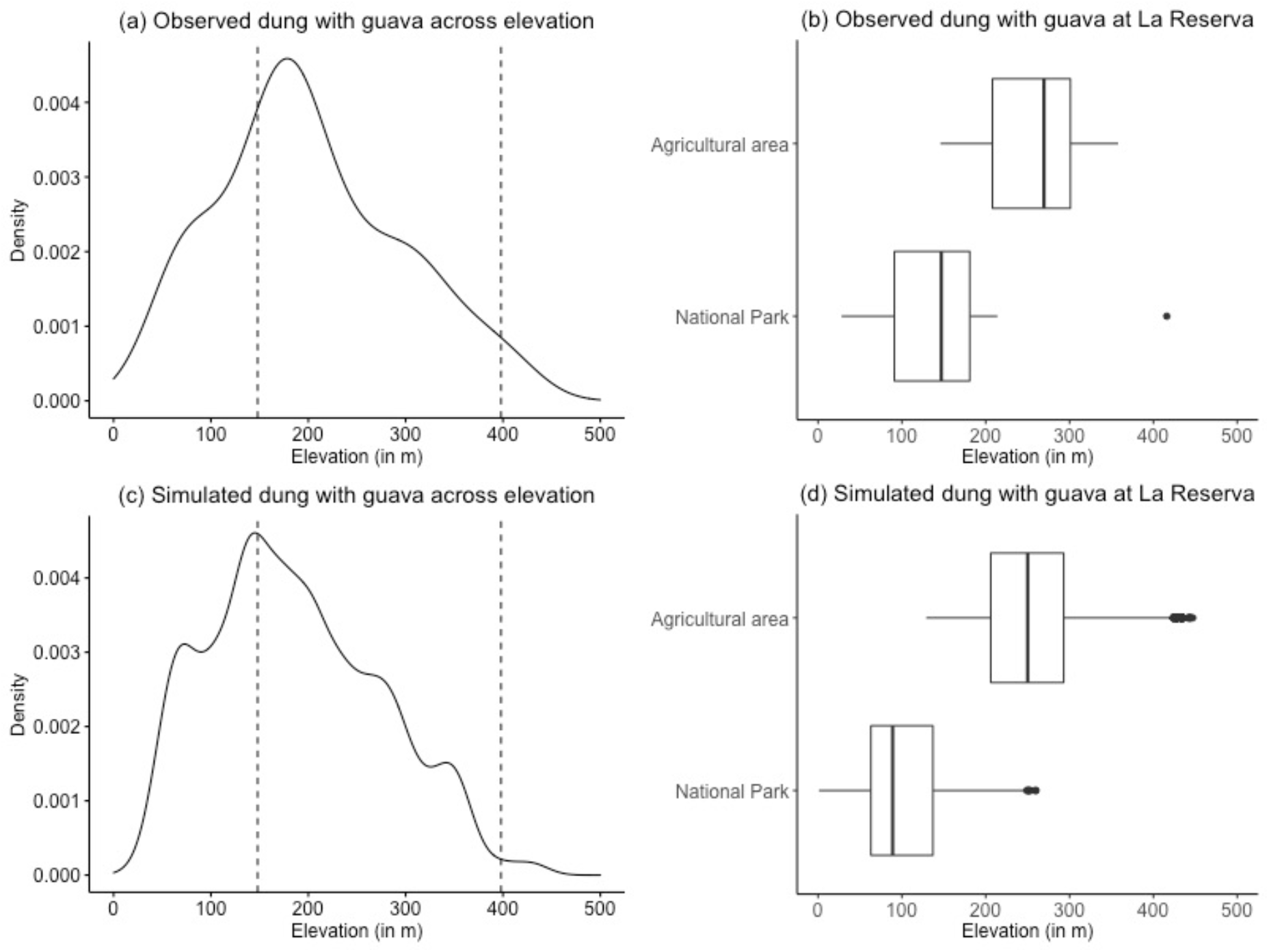
Model simulation captures empirical dispersal of guava seeds by migrating tortoises. Distribution of observed and simulated tortoise dung piles with guava across the elevational gradient of Santa Cruz Island. Observed (a) and simulated (c) guava seed deposition (left panel density plots) and location in agricultural- and national park areas deposition (b, d, right panel boxplots) across the elevational gradient of site “La Reserva” in Santa Cruz, Galapagos.

### 3.2 Estimating seed fate using landcover and species distribution models

By combining the guava seed rain simulated from our model (Fig. 2b) with landcover categories, we provide a spatial and ecological context in which guava seeds were deposited. Simulated guava seeds were deposited in five different landcover classes, with a pattern of deposition in these different landcover classes that largely matches the one from empirical data (Figs. 3b,d and S4). Guava seeds were deposited mostly in agricultural areas but also reached the Galapagos National Park (Fig. 3b,d). Some discrepancies exist between simulation and empirical data, notably that the model predicts guava seeds to be deposited substantially more than suggested from empirical data in areas occupied by invasive species but less than empirical data suggests in evergreen forests and shrubs (Fig. S4). In addition, when applying our threshold guava SDM mask, we were able to create a spatially explicit prediction of successful seed dispersal for *C. porteri*, simulating guava seeds with the potential to germinate and establish under current climatic conditions (Fig. S5).

## 4.0 Discussion

In this study, we developed a novel modeling framework for simulating the dispersal and fate of propagules carried by migratory animals. Our model couples empirical data on animal behavior related to feeding and movement as well as data on the geographical distribution and climatic/habitat suitability of the dispersed species. Our model is data-driven, spatially explicit, and it accounts for intraspecific variation in animal migratory movements. It provides a quantitative framework for predicting long-distance propagule dispersal events by animals, which have rarely been quantitatively predicted despite their ecological importance (Nathan & Muller-Landau 2000). In particular, our model is able to predict the magnitude and direction under which a migratory animal population disperses plant species into novel habitats, and quantify how the seasonal migration of herbivores may be a major seed dispersal vector and expand the distribution range of invasive plant species.

We applied our modeling framework to guava seed dispersal by Galapagos tortoises (*C. porteri*) on Santa Cruz Island, which is a data-rich system with relevance for conservation. Our model has good predictive ability for the pattern of spread of guava seeds by migrating tortoises (Fig. 3), and can therefore be used for making predictions beyond the individuals and areas where data were collected. We predicted clusters of heavy seed dispersal alongside tortoises’ downslope migration range (Fig. 2) and identified areas most likely to see germination success and establishment (Fig. S5). We found that Galapagos tortoises’ seed dispersal may particularly contribute to guava downslope range expansion on Santa Cruz Island, due to both long gut retention time and tortoise’s long-distance migration across vegetation zones. In particular, we predict that tortoises are dispersing a significant amount of guava seeds into the Galapagos National Park, which has important consequences for the native flora as guava is an invasive species that has already altered natural ecosystems on the Galapagos (Wiggins & Porter 1971; Weber 2003) and is threatening local and endemic plant species (de Lourdes Torres & Mena 2018). Our results also highlight that the frequency at which behavior (i.e. migratory movements, eating, excreting) is repeated during the migration season has an important role to play in predicting long-distance dispersal events (Fig. S3), thus highlighting an avenue for further research.

Our framework is flexible and modular as it allows for increasing complexity and the integration of multiple data sources (Fig. 1). For example, if the number of seeds excreted and germination success are unknown for a study system, this information could be omitted and the model could simply predict the location of seed dispersal events across space. More information on population distribution and individual-level movement, which is increasing exponentially thanks to advances in citizen science and tracking technology (Kays *et al*. 2020), could be used to calibrate more accurately the animal migration module (i.e. model components UD, ORSIM and eRTG; Fig. 1). In addition, besides using SDMs, other proxies for seed fate such as additional landcover information or microclimate based on fine scale topography could be employed (Leempoel *et al*. 2015; Maclean *et al*. 2019; van Toor *et al*. 2019). *In situ* germination trials of seeds along environmental gradients would provide greatest insights in the seed dispersal efficiency of a migratory animal species, which is relevant as plant mortality depends on environmental conditions and is highest when plants are seedlings (Terborgh *et al*. 2014). Future studies could also include plant demography in addition to the movement ecology of seeds in order to fully model the contribution of animal-mediated seed dispersal to the range expansion of an invasive species (Beckman *et al*. 2019).

A research avenue for which our modeling framework could provide an important contribution is investigating the impact of environmental change on ecosystem processes. In particular, change in climate and habitat quality might alter animal migration patterns and seeds germination success, thus affecting animal-mediated seed dispersal. As it integrates models of animal movement influenced by the distribution of resources (using NDVI here but other data could be used) and seeds germination success based on climate suitability from species distribution modeling (Ellis-Soto et al. 2017), our approach could be used to make future predictions for where will seeds disperse and successfully establish under various scenarios of environmental change. Thus, the predictive ability and flexibility of our framework makes it a valuable tool for investigating seed dispersal under climate change, which cannot be easily predicted using existing modeling approaches.

Our framework can be applied to a wide range of systems where ecosystem processes are affected by migratory animal species. It would be interesting for example to investigate how migratory populations living in different environmental settings would spread an invasive plant in different ways, such as for example how different populations of tortoises spread guava on different islands in the Galapagos. Such predictions would allow identifying which islands and ecosystems are most at risk of invasion by guava and it would therefore inform where to focus conservation efforts. In addition, our modeling framework could be used to investigate the relative roles of different animal species, given their movement ability and patterns, in spreading the seeds of an invasive plant throughout an ecosystem, which is a promising research avenue with important consequences for conservation and ecosystem functioning. In the case study presented here, giant tortoises are not the only animals capable of dispersing seeds across Galapagos Islands. Galapagos mockingbirds (*Mimus parvulus*), Darwin finches (Geospiza spp.) and introduced cattle, pigs and goats also consume guava, but they move shorter distances (Buddenhagen & Jewell 2006) and cross less from agricultural areas in the highlands to the national park in the lowlands when compared to tortoises. Humans in the Galapagos are also potential long-distance dispersers of guava (Auffret *et al*. 2014). It would be informative to use our model to quantify the relative contributions of these different vectors to the spread of guava. Finally, our modeling framework could also be used to investigate ecosystem processes other than seed dispersal. For example, it would be possible to apply it to quantitatively model how migratory animals affect nutrient transport across ecosystems, which can be quantified either through empirical measurements of excretion rate or metabolic measurements. These different applications can improve our understanding on how animals connect ecosystems and landscapes across spatiotemporal scales and have important conservation implications for the management of ecosystems that include migratory animals.

## 5.0 Acknowledgements

We would like to thank Stephen Blake and Guillaume Bastille-Rousseau for the feedback on the manuscript, as well as Kamran Safi for sharing the eRTG code. We also thank Freddy Cabrera for collection of tortoise telemetry data. We thank Patricia Jaramillo for mentoring of D.E.S in guava seed identification. D.E.S. acknowledges support from the Rufford Foundation. We thank the Galapagos National Park, and the Charles Darwin Foundation for providing critical logistical, administrative and technical support facilitating data collection and providing permits.

## 7.0 Data Availability Statement

Land cover classification maps are available on the supplementary material of Rivas-Torres *et al*. (2018) and guava distribution in Santa Cruz Island is available on the supplementary material of Laso *et al*. (2019). Galápagos giant tortoise tracking data used in this study will be archived in a public Movebank data repository, should the manuscript be accepted, and the data DOI will be included at the end of the article. The computer code used for this study is available at https://github.com/msomveille/galapagos-tortoises.git.

## Notes

### Competing Interest Statement

The authors have declared no competing interest.

## References

Auffret, A.G., Berg, J. & Cousins, S.A.O. (2014). The geography of human-mediated dispersal. Divers. Distrib., 20, 1450–1456.

Bastille-Rousseau, G., Yackulic, C., Gibbs, J., Frair, J., Cabrera, F. & Blake, S. (2019). Migration triggers in a large herbivore: Galapagos giant tortoises navigating resource gradients on volcanoes. Ecology, 100, e02658.

Bauer, S. & Hoye, B.J. (2014). Migratory animals couple biodiversity and ecosystem functioning worldwide. Science, 344, 1242552.

Beckman, N.G., Aslan, C.E., Rogers, H.S., Kogan, O., Bronstein, J.L., Bullock, J.M., et al. (2019). Advancing an interdisciplinary framework to study seed dispersal ecology. AoB Plants, 12, plz048.

Benitez-Capistros, F., Camperio, G., Hugé, J., Dahdouh-Guebas, F. & Koedam, N. (2018). Emergent conservation conflicts in the galapagos islands: Human-giant tortoise interactions in the rural area of Santa Cruz Island. PLoS One, 13, e0202268.

Benitez-Capistros, F., Couenberg, P., Nieto, A., Cabrera, F. & Blake, S. (2019). Identifying Shared Strategies and Solutions to the Human–Giant Tortoise Interactions in Santa Cruz, Galapagos: A Nominal Group Technique Application. Sustainability, 11, 2937.

Blake, S., Guézou, A., Deem, S.L., Yackulic, C.B. & Cabrera, F. (2015). The Dominance of Introduced Plant Species in the Diets of Migratory Galapagos Tortoises Increases with Elevation on a Human-Occupied Island. Biotropica, 47, 246–258.

Blake, S., Wikelski, M., Cabrera, F., Guezou, A., Silva, M., Sadeghayobi, E., et al. (2012). Seed dispersal by Galapagos tortoises. J. Biogeogr., 39, 1961–1972.

Blake, S., Yackulic, C.B., Cabrera, F., Tapia, W., Gibbs, J.P., Kümmeth, F., et al. (2013). Vegetation dynamics drive segregation by body size in Galapagos tortoises migrating across altitudinal gradients. J. Anim. Ecol., 82, 310–321.

Buddenhagen, C. & Jewell, K.J. (2006). Invasive plant seed viability after processing by some endemic Galapagos birds. Ornitol. Neotrop., 17, 73–80.

Bullock, J.M., Mallada González, L., Tamme, R., Götzenberger, L., White, S.M., Pärtel, M., et al. (2017). A synthesis of empirical plant dispersal kernels. J. Ecol.

Calenge, C. (2006). The package “adehabitat” for the R software: A tool for the analysis of space and habitat use by animals. Ecol. Modell., 197, 516–519.

CGREG (2015). Censo de Unidades de Producción Agropecuaria de Galápagos 2014 (UPA), Consejo de Gobierno del Régimen Especial de Galápagos.

Côrtes, M.C. & Uriarte, M. (2013). Integrating frugivory and animal movement: a review of the evidence and implications for scaling seed dispersal. Biol. Rev., 88, 255–272.

Dokter, A.M., Farnsworth, A., Fink, D., Ruiz-Gutierrez, V., Hochachka, W.M., La Sorte, F.A., et al. (2018). Seasonal abundance and survival of North America’s migratory avifauna determined by weather radar. Nat. Ecol. Evol., 2, 1603–1609.

Earl, J.E. & Zollner, P.A. (2017). Advancing research on animal-transported subsidies by integrating animal movement and ecosystem modelling. J. Anim. Ecol., 86, 987–997.

Ellis-Soto, D. (2020). Giant tortoises connecting terrestrial and freshwater ecosystems in Santa Cruz Island. In: Galapagos Giant Tortoises (eds. Gibbs, J.P., Cayot, L.J. & Tapia, W.). Elsevier, Amsterdam, p. 286.

Ellis-Soto, D., Blake, S., Soultan, A., Guézou, A., Cabrera, F. & Lötters, S. (2017). Plant species dispersed by Galapagos tortoises surf the wave of habitat suitability under anthropogenic climate change. PLoS One, 12, e0181333.

Ellis-Soto, D., Ferraro, K.M., Rizzuto, M., Briggs, E., Monk, J.D. & Schmitz, O.J. (2020). A methodological roadmap to quantify animal-vectored spatial ecosystem subsidies. EcoEvoRxiv.

Fieberg, J. & Kochanny, C.O. (2005). Quanitfying home-range overlap: the importance of the utilization distribution. J. Wildl. Manage., 69, 1346–1359.

Gardener, M.R., Atkinson, R. & Renteria, J.L. (2010). Eradications and people: lessons from the plant eradication program in Galapagos. Restoration Ecology, 18, 20–29.

Gibbs, J.P., Sterling, E.J. & Zabala, F.J. (2010). Giant tortoises as ecological engineers: A long-term quasi-experiment in the Galapagos Islands. Biotropica, 42, 208–214.

Guézou, A., Trueman, M., Buddenhagen, C.E., Chamorro, S., Guerrero, A.M., Pozo, P., et al. (2010). An extensive alien plant inventory from the inhabited areas of galapagos. PLoS One, 5, e10276.

Hawkes, L.A., Balachandran, S., Batbayar, N., Butler, P.J., Frappell, P.B., Milsom, W.K., et al. (2011). The trans-Himalayan flights of bar-headed geese (Anser indicus). Proc. Natl. Acad. Sci., 108, 9516–9519.

Hijmans, R.J., Cameron, S.E., Parra, J.L., Jones, P.G. & Jarvis, A. (2005). Very high resolution interpolated climate surfaces for global land areas. Int. J. Climatol., 25, 1965–1978

Hillier, F.S. & Lieberman, G.J. (1990). Introduction to mathematical programming. McGraw-Hill, New York, NY.

Hu, G., Chapman, J.W., Lim, K.S., Reynolds, D.R., Clark, S.J., Horvitz, N., et al. (2016). Mass seasonal bioflows of high-flying insect migrants. Science, 354, 1584–1587.

Jesmer, B.R., Merkle, J.A., Goheen, J.R., Aikens, E.O., Beck, J.L., Courtemanch, A.B., et al. (2018). Is ungulate migration culturally transmitted? Evidence of social learning from translocated animals. Science, 361, 1023–1025.

Kays, R., Crofoot, M.C., Jetz, W. & Wikelski, M. (2015). Terrestrial animal tracking as an eye on life and planet. Science, 348, aaa2478.

Kays, R., McShea, W.J. & Wikelski, M. (2020). Born-digital biodiversity data: millions and billions. Divers. Distrib., 26, 644–648.

Kendrick, G.A., Waycott, M., Carruthers, T.J.B., Cambridge, M.L., Hovey, R., Krauss, S.L., et al. (2012). The Central Role of Dispersal in the Maintenance and Persistence of Seagrass Populations. Bioscience, 62, 56–65.

Kleyheeg, E., Fiedler, W., Safi, K., Waldenström, J., Wikelski, M. & van Toor, M.L. (2019). A Comprehensive Model for the Quantitative Estimation of Seed Dispersal by Migratory Mallards. Front. Ecol. Evol., 7, 1–14.

Kleyheeg, E., Treep, J., de Jager, M., Nolet, B.A. & Soons, M.B. (2017). Seed dispersal distributions resulting from landscape-dependent daily movement behaviour of a key vector species, *Anas platyrhynchos*. J. Ecol., 105, 1279–1289

Laso, F.J., Ben, L., Rivas-torres, G., Sampedro, C. & Arce-nazario, J. (2019). Land Cover Classification of Complex Agroecosystems in the Non-Protected Highlands of the Galapagos Islands. Remote Sens., 12, 65.

Leempoel, K., Parisod, C., Geiser, C., Daprà, L., Vittoz, P. & Joost, S. (2015). Very high-resolution digital elevation models: are multi-scale derived variables ecologically relevant? Methods Ecol. Evol., 6, 1373–1383.

Loreau, M., Mouquet, N. & Holt, R.D. (2003). Meta-ecosystems: a theoretical framework for a spatial ecosystem ecology. Ecol. Let., 6, 673–679.

de Lourdes Torres, M. & Mena, C.F. (2018). Understanding Invasive Species in the Galapagos Islands. Springer.

Maclean, I.M.D., Mosedale, J.R. & Bennie, J.J. (2019). Microclima: An r package for modelling meso- and microclimate. Methods Ecol. Evol., 10, 280–290.

Malhi, Y., Doughty, C.E., Galetti, M., Smith, F.A., Svenning, J.-C. & Terborgh, J.W. (2016). Megafauna and ecosystem function from the Pleistocene to the Anthropocene. Proc. Natl. Acad. Sci., 113, 838–846.

Nathan, R. (2006). Long-distance dispersal of plants. Science, 313, 786–788.

Nathan, R., Getz, W.M., Revilla, E., Holyoak, M., Kadmon, R., Saltz, D., et al. (2008). A movement ecology paradigm for unifying organismal movement research Ran. Proc. Natl. Acad. Sci., 105, 19052–19059.

Nathan, R., Klein, E., Robledo-Arnuncio, J.J. & Revilla, E. (2012). Dispersal kernels: review, Dispersal Ecology and Evolution. Oxford University Press.

Nathan, R. & Muller-Landau, H.C. (2000). Spatial patterns of seed dispersal, their determinants and consequences for recruitment. Trends Ecol. Evol., 15, 278–285.

Neuschulz, E.L., Mueller, T., Schleuning, M. & Böhning-Gaese, K. (2016). Pollination and seed dispersal are the most threatened processes of plant regeneration. Sci. Rep., 6, 6–11.

Oleksy, R., Giuggioli, L., McKetterick, T.J., Racey, P.A. & Jones, G. (2017). Flying foxes create extensive seed shadows and enhance germination success of pioneer plant species in deforested Madagascan landscapes. PLoS One, 12, 1–17.

Pele, O. & Werman, M. (2008). A linear time histogram metric for improved SIFT matching. Computer Vision – ECCV 2008, Marseille, France, 495–508.

Pele, O. & Werman, M. (2009). Fast and robust earth mover’s distances. Proc. 2009 IEEE 12^th^ Int. Conf. on Computer Vision, Kyoto, Japan, 460–467.

Pires, M.M., Guimarães, P.R., Galetti, M. & Jordano, P. (2018). Pleistocene megafaunal extinctions and the functional loss of long-distance seed-dispersal services. Ecography, 41, 153–163.

Rivas-Torres, G.F., Benítez, F.L., Rueda, D., Sevilla, C. & Mena, C.F. (2018a). A methodology for mapping native and invasive vegetation coverage in archipelagos: An example from the Galápagos Islands. Prog. Phys. Geogr.

Rivas-Torres, G.F., Benítez, F.L., Rueda, D., Sevilla, C. & Mena, C.F. (2018b). A methodology for mapping native and invasive vegetation coverage in archipelagos: An example from the Galápagos Islands. Prog. Phys. Geogr. Earth Environ., 42, 83–111.

Sadeghayobi, E., Blake, S., Wikelski, M., Gibbs, J., Mackie, R. & Cabrera, F. (2011). Comparative Biochemistry and Physiology, Part A Digesta retention time in the Galápagos tortoise (*Chelonoidis nigra*). Comp. Biochem. Physiol. Part A, 160, 493–497.

Schmitz, O.J., Wilmers, C.C., Leroux, S.J., Doughty, C.E., Atwood, T.B., Galetti, M., et al. (2018). Animals and the zoogeochemistry of the carbon cycle. Science, 362, eaar3213.

Singh, N.J., Allen, A.M. & Ericsson, G. (2016). Quantifying Migration Behaviour Using Net Squared Displacement Approach: Clarifications and Caveats. PLoS One, 11, 1–20.

Smouse, P.E., Focardi, S., Moorcroft, P.R., Kie, J.G., Forester, J.D. & Morales, J.M. (2010). Stochastic modelling of animal movement. Philos. Trans. R. Soc. B Biol. Sci., 365, 2201–2211.

Subalusky, A.L. & Post, D.M. (2018). Context dependency of animal resource subsidies. Biol. Rev., 94, 517–538.

Technitis, G., Othman, W., Safi, K. & Weibel, R. (2015). From A to B, randomly: a point-to-point random trajectory generator for animal movement. Int. J. Geogr. Inf. Sci., 29, 912–934.

Terborgh, J., Zhu, K., Alvarez-Loayza, P. & Cornejo Valverde, F. (2014). How many seeds does it take to make a sapling? Ecology.

van Toor, M.L., O’Mara, M.T., Abedi-Lartey, M., Wikelski, M., Fahr, J. & Dechmann, D.K.N. (2019). Linking colony size with quantitative estimates of ecosystem services of African fruit bats. Curr. Biol., 29, R237–R238.

Travis, J.M.J., Delgado, M., Bocedi, G., Baguette, M., Bartoń, K., Bonte, D., et al. (2013). Dispersal and species’ responses to climate change. Oikos, 122, 1532–1540.

Trueman, M., Standish, R.J. & Hobbs, R.J. (2014). Identifying management options for modified vegetation: Application of the novel ecosystems framework to a case study in the Galapagos Islands. Biol. Conserv., 172, 37–48.

Tucker, M.A., Busana, M., Huijbregts, M.A. & Ford, A.T. (2021). Human-induced reduction in mammalian movements impacts seed dispersal in the tropics. Ecography DOI: doi.org/10.111/ecog.05210.

Walsh, S.J., McCleary, A.L., Mena, C.F., Shao, Y., Tuttle, J.P., González, A., et al. (2008). QuickBird and Hyperion data analysis of an invasive plant species in the Galapagos Islands of Ecuador: Implications for control and land use management. Remote Sens. Environ., 112, 1927–1941.

Weber, E. (2003). Invasive Plant Species of the World: A Reference Guide to Environmental Weeds. illustrate. CABI Publishing Series.

Wiggins, I.L. & Porter, D.M. (1971). Flora of the Galapagos Islands. Stanford University Press, Stanford.

Wilcove, D.S. & Wikelski, M. (2008). Going, Going, Gone: Is Animal Migration Disappearing. PLoS Biol., 6, e188.

Yackulic, C.B., Blake, S. & Bastille-Rousseau, G. (2016). Benefits of the destinations, not costs of the journeys, shape partial migration patterns in Galapagos tortoises. J. Anim. Ecol., 86, 972–982.

Zwolak, R. (2018). How intraspecific variation in seed-dispersing animals matters for plants. Biol. Rev., 93, 897–913.

