## Supplementary Material for "Linking animal migration and ecosystem processes: data-driven simulation of propagule dispersal by migratory herbivores"

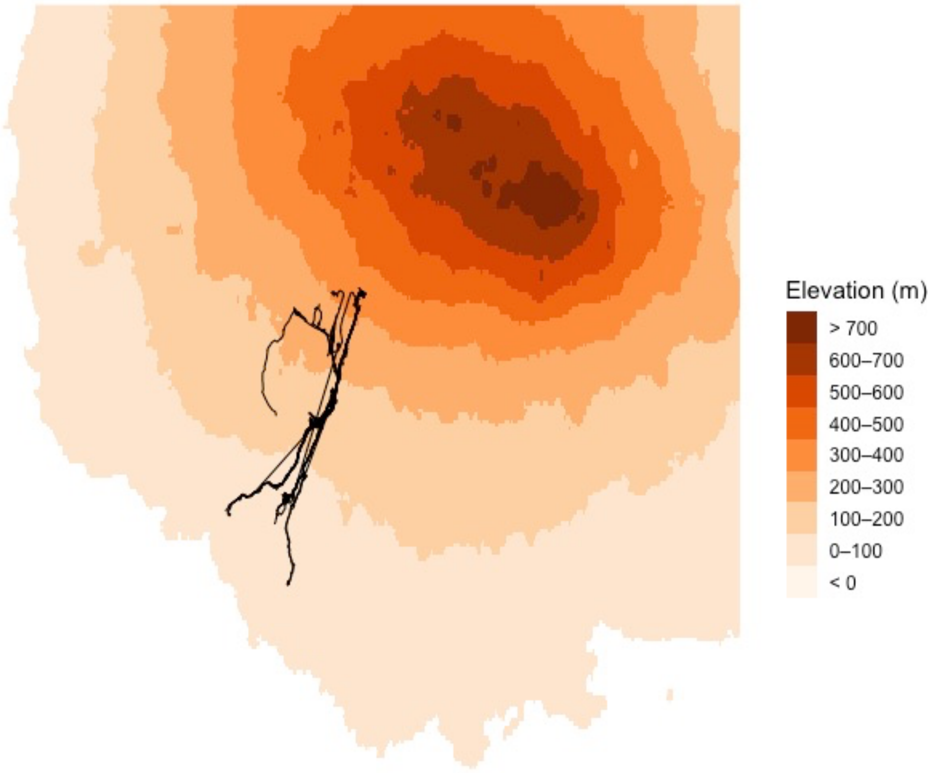


**Figure S1. Empirical tracks of migrating tortoises used to calibrate the model**. 19 migratory tracks, belonging to 5 individuals, are represented.

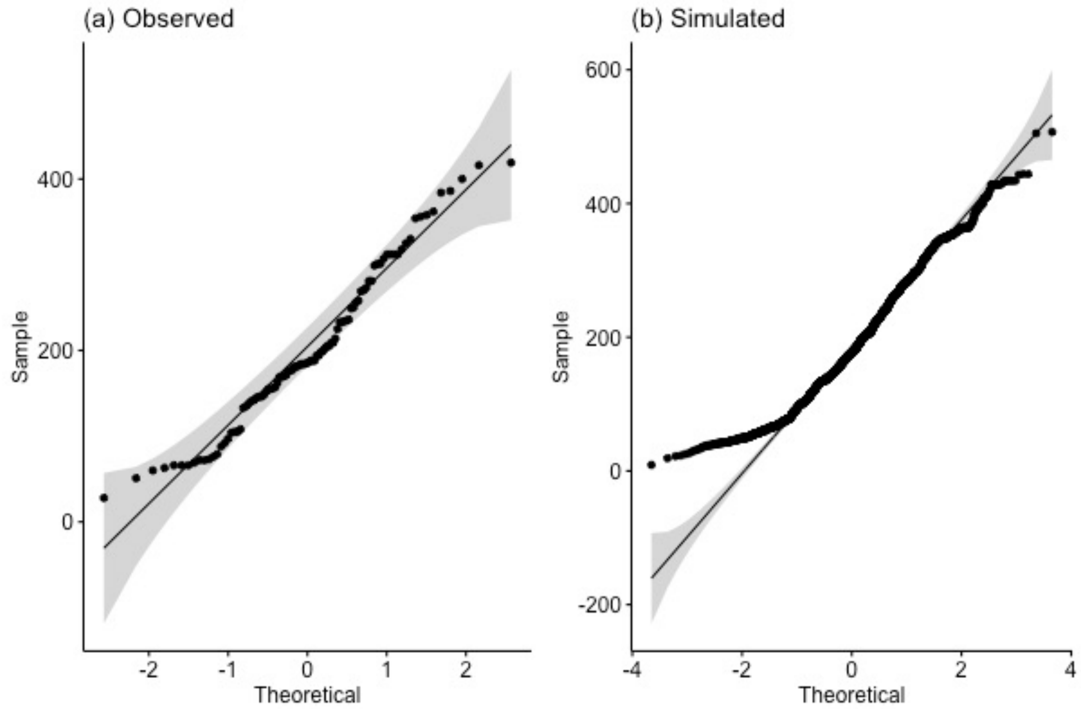


**Figure S2.** Quantile-Quantile plot of observed (a) and simulated (b) dung piles with guava across elevation in Santa Cruz Island

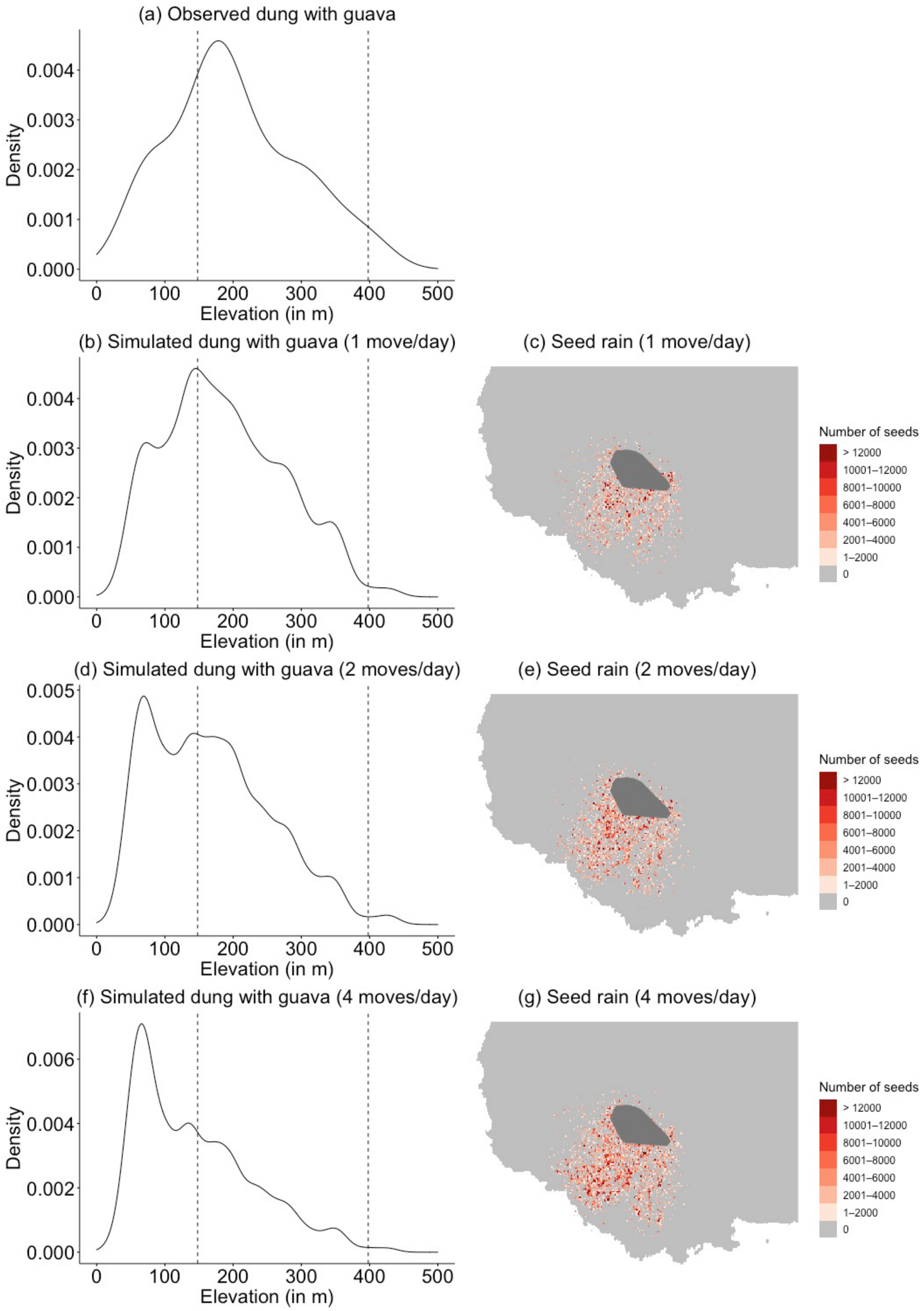

(a) Observed dung with guava
Density
Elevation (in m)
(b) Simulated dung with guava (1 move/day)
(c) Seed rain (1 move/day)
(d) Simulated dung with guava (2 moves/day)
(e) Seed rain (2 moves/day)
(f) Simulated dung with guava (4 moves/day)
(g) Seed rain (4 moves/day)
Number of seeds
> 12000
10001–12000
8001–10000
6001–8000
4001–6000
2001–4000
1–2000
0

**Figure S3. Sensitivity analysis for different frequency of migratory movements.** Density of (a) observed and (b,d,f) simulated tortoise dung piles containing guava seeds across the elevational gradient of Santa Cruz Island, with the associated predicted seed rain (c,e,g). These results are obtained for simulated migrating tortoises engaging in migratory movement (b–c) once a day, (d–e) twice a day, and (f–g) four times a day. In dark grey (c,e,g): highland distribution of the population.

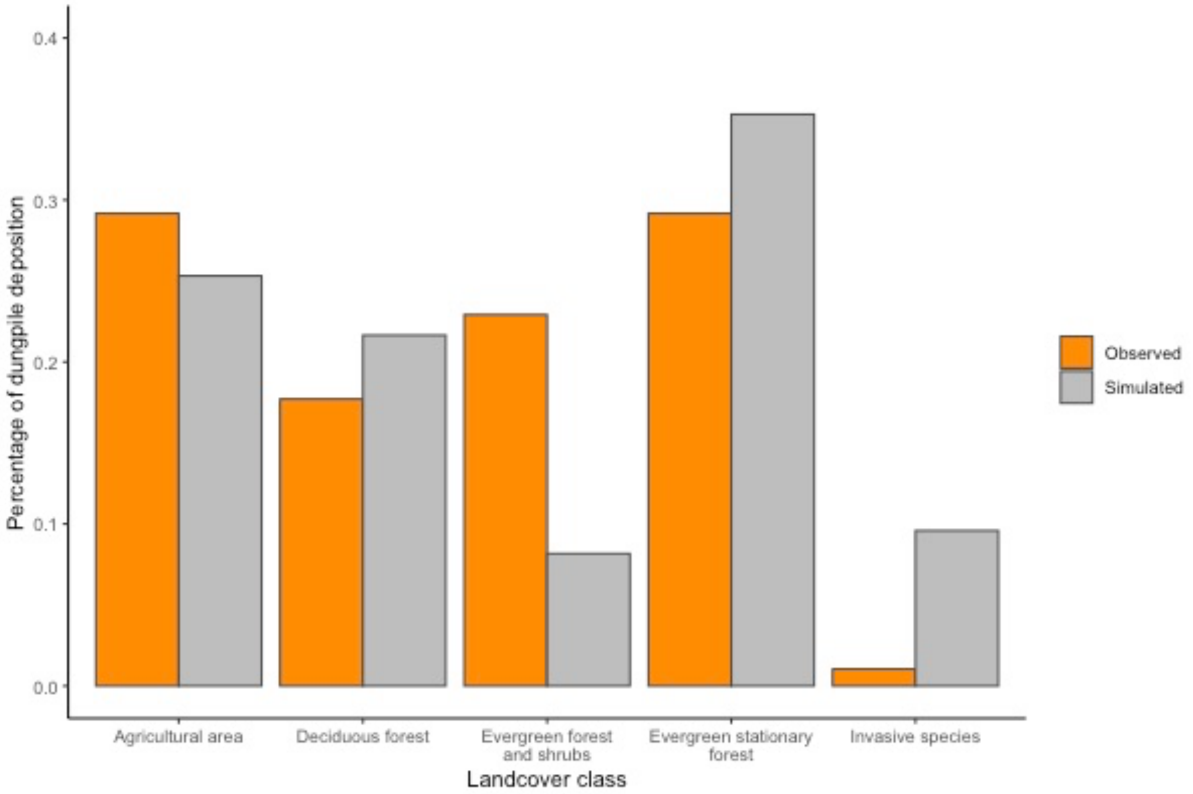


**Figure S4. Landcover types under which simulated guava seeds are deposited.** Spatial deposition of observed (orange) and simulated (grey) dung piles containing guava for different landcover classes from Rivas-Torres *et al.* (2018a).

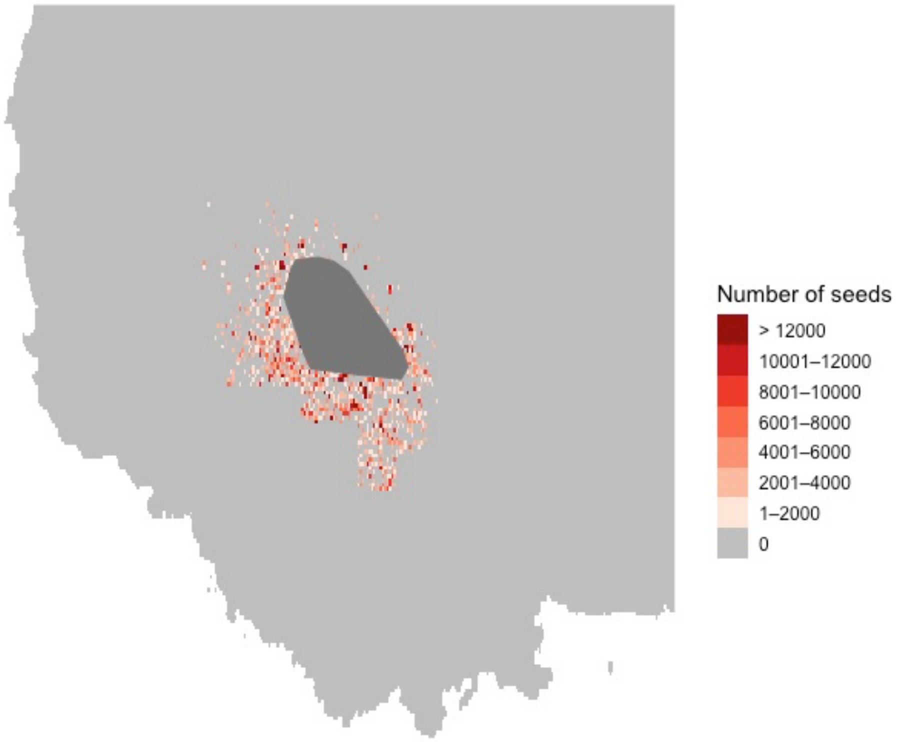


**Figure S5. Estimated successful seed dispersal.** Density of germinated guava seeds dispersed by simulated migrating tortoises obtained by combining the simulated seed rain with the germination success of guava seeds based on a species distribution model (Ellis-Soto *et al.* 2017). In dark grey: highland distribution of the population.
